# CRISPR-Cas9 modified bacteriophage for treatment of *Staphylococcus aureus* induced osteomyelitis and soft tissue infection

**DOI:** 10.1101/705830

**Authors:** Leah K. Horstemeyer, JooYoun Park, Elizabeth A. Swanson, Mary Catherine Beard, Emily M. McCabe, Anna S. Rourke, Keun Seok Seo, Alicia K. Olivier, Lauren B. Priddy

**Affiliations:** Department of Agricultural and Biological Engineering, Mississippi State University, Mississippi State, MS, USA; Department of Basic Sciences, College of Veterinary Medicine, Mississippi State University, Mississippi State, MS, USA; Department of Clinical Sciences, College of Veterinary Medicine, Mississippi State University, Mississippi State, MS, USA; Department of Pathobiology and Population Medicine, College of Veterinary Medicine, Mississippi State University, Mississippi State, MS, USA

## Abstract

Osteomyelitis, or bone infection, is often induced by antibiotic resistant *Staphylococcus aureus* strains of bacteria. Although debridement and long-term administration of antibiotics are the gold standard for osteomyelitis treatment, the increase in prevalence of antibiotic resistant bacterial strains limits the ability of clinicians to effectively treat infection. Bacteriophages (phages), viruses that effectively lyse bacteria, have gained recent attention for their high specificity, non-toxicity, and the low likelihood of resistance development by pathogens. Previously, we have shown that CRISPR-Cas9 genomic editing techniques could be utilized to expand bacteriophage host range and enhance bactericidal activity through modification of the tail fiber protein, as well as improve safety with removal of major virulence genes. In a dermal infection study, these CRISPR-Cas9 phages reduced bacterial load relative to unmodified phage. Thus, we hypothesized this bacteriophage would be effective to mitigate infection from a biofilm forming *S. aureus* strain *in vitro* and *in vivo*. *In vitro*, qualitative fluorescent imaging demonstrated superiority of phage to conventional vancomycin and fosfomycin antibiotics against *S. aureus* biofilm. Quantitative antibiofilm effects increased over time for fosfomycin, phage, and fosfomycin-phage (dual) therapeutics delivered via alginate hydrogel. We developed an *in vivo* rat model of osteomyelitis and soft tissue infection that was reproducible and challenging and enabled longitudinal monitoring of infection progression. Using this model, phage (with and without fosfomycin) delivered via alginate hydrogel were successful in reducing soft tissue infection but not bone infection, based on bacteriological, histological, and scanning electron microscopy analyses. Notably, the efficacy of phage at mitigating soft tissue infection was equal to that of high dose fosfomycin. Future research may utilize this model as a platform for evaluation of therapeutic type and dose, and alternate delivery vehicles for osteomyelitis mitigation.

## Introduction

For nearly a century, antibiotics have been a vital resource utilized by clinicians to eliminate infection, with nearly 270 million prescriptions dispensed in 2015 alone.[1] Antibiotics are utilized for a variety of infections, from common otitis externa (“swimmers ear”) to severe endocarditis, pneumonia, meningitis or osteomyelitis. Although antibiotics are typically able to clear infection, antibiotic resistant strains of bacteria continue to emerge. It is not as lucrative, nor as feasible, for pharmaceutical companies to develop novel antibiotics at the rates that these multi-drug resistant (MDR) bacterial strains are isolated. Nationally, approximately $2.2 billion is spent annually to treat MDR bacterial infections.[2] By 2050, it is estimated that nearly 10 million people could die each year due to resistant strains of bacteria.[3]

*Staphylococcus aureus* (*S. aureus*), a gram-positive bacterial strain, is one of the most commonly isolated and arguably one of the most detrimental pathogens with antibiotic resistance. One of the most common antibiotic resistant strains of *S. aureus* is methicillin-resistant *S. aureus* (MRSA). MRSA alone was responsible for over 80,000 reported infections in 2011 alone, of which 11,285 resulted in death.[4] *S. aureus* is able to achieve antibiotic resistance with genomic changes such as altered synthesis of peptidoglycan, a major component of the bacterial cell wall. Additionally, some strains of *S. aureus* can produce biofilms, an extracellular polymeric matrix including dead bacterial cells, which surrounds and protects the living, underlying layer of *S. aureus*.[5] These biofilms can be difficult to penetrate, and oftentimes require surgical intervention to remove.

Difficulties in treating osteomyelitis, or the infection of bone, have been exacerbated by the rise of antibiotic resistant bacterial strains, particularly *S. aureus* strains, which are the most common cause of bone infection.[6] Of diabetic foot ulcers, which occur in 25% of diabetic patients, approximately 20% will spread to nearby bony hosts and result in osteomyelitis.[7] As diabetic diagnoses continue to increase in the United States with an expected 55 million to be afflicted by 2030, osteomyelitis infections will be an ongoing challenge for the healthcare community.[8] It is essential that new therapeutics be engineered and tested, for rapid translation into clinical use.

Bacteriophages (phages), or viruses that kill their bacterial hosts, are one class of therapeutics that have gained attention in recent years due to their high specificity, non-toxicity, and abundancy in nature.[9,10] Phages have been used for decades in Eastern Europe but have not yet been adopted in the United States or other countries. This may be due to public concern regarding elective viral use, issues concerning commercial phage production, and/or the ability to fund and validate clinical trials.[11] Nonetheless, the potential benefits of this treatment have been indicated by results of clinical trials of phages for treating diabetic foot ulcers, chronic otitis, and urinary tract infections[11–13]. In April 2019, data from clinical trials were published from Sydney, Australia, where intravenous (IV) administration of phage was utilized for *Staphylococcus* infection treatment. Marked reduction of *staphylococci* with no adverse events were reported.[14] In the United States as of January 2019, IV administration of phage for ventricular assist device infection treatment received approval for phase I/II clinical trials.[15] Collectively, these clinical trials demonstrate the efficacy of bacteriophage therapeutics and suggest their potential utility against MDR bacterial strains.

Hydrogels are a commonly used, easily tailored delivery vehicle for therapeutics for a wide variety of ailments, including osteomyelitis[6,16,17]. Alginate hydrogels are injectable, well characterized, and biocompatible.[18,19] Furthermore, bacteriophages have been successfully delivered to sites of infection with various hydrogel-based delivery systems in previous studies.[16,17,20]

Although the high specificity of phages can be beneficial for treating a known, single species, specificity of these viruses can make polymicrobial infection mitigation challenging. In the clinical scenario, it is ideal for health care providers to administer one broad-spectrum drug immediately upon patient presentation, rather than spend time identifying the causative agents of infection. Previously, we have utilized CRISPR-Cas9 to modify temperate bacteriophage, which effectively removed major virulence genes and expanded host range via modifications to the tail fiber protein (which codes host specificity). *In vitro* testing revealed the improvements of bacteriophage bactericidal activity due to this CRISPR-Cas9 system.[21]Within 6h of treatment, the CRISPR-Cas9 phage effectively killed 1×10^5^ CFU *S. aureus* culture. With native, unmodified phage treatment, the culture was found to increase to approximately 1×10^9^ CFU. Similar effects were noted in an *in vivo* dermal infection study, where CRISPR-Cas9 phage treatment resulted in nearly complete mitigation of dermal infections (~1 log CFU/g tissue), while treatment with unmodified phage resulted in a significantly higher bacterial load (~3.5 log CFU/g tissue).[21]

The objectives of the present work were: (i) to develop a green fluorescent protein (GFP) integrated *S. aureus* strain (ATCC 6538-GFP), modify bacteriophage using CRISPR-Cas9, and evaluate the bactericidal efficacy of our CRISPR-Cas9 modified bacteriophage *in vitro*, compared to conventional antibiotics, and (ii) to develop an *in vivo* model of osteomyelitis and soft tissue infection using this biofilm forming *S. aureus* strain, and use it to assess the antimicrobial effects of bacteriophage, antibiotic, and dual bacteriophage-antibiotic therapies via histological, radiographic, and bacteriological analyses. Our hypothesis was that CRISPR-Cas9 modified bacteriophage would be effective against *S. aureus* infection *in vitro* and in the femur and contiguous soft tissue *in vivo*.

## Materials and Methods

### Bacterial strains and culture

For a stable quantification of biofilm, *S. aureus* strain ATCC 6538 was genetically modified to contain chromosomally integrated green fluorescent protein (GFP), as previously described.[22] Briefly, *S. aureus* strain ATCC 6538 was transduced with a temperature sensitive plasmid pTH100 harboring the GFP gene by electroporation and cultured in a brain heart infusion (BHI) agar plate supplemented with chloramphenicol (BHI-CM) at 30°C, a plasmid replication permissive temperature. To promote the first homologous recombination and cure pTH100, a single colony grown in a BHI-CM plate was transferred to a fresh BHI-CM plate and cultured at 42°C, a plasmid replication non-permissive temperature. To promote the second homologous recombination, which removed the plasmid and resulted in a loss of chloramphenicol resistance but maintained the GFP phenotype, a single colony was inoculated into BHI broth and cultured at 37°C overnight. A serial dilution of culture was inoculated onto a BHI plate and incubated at 37°C overnight. A GFP positive single colony checked by ultraviolet lamp was randomly selected and streaked onto BHI and BHI-CM. A colony that was both GFP positive and sensitive to chloramphenicol, indicating the integration of the GFP gene into the chromosome and removal of plasmid, was selected for experiments (ATCC 6538-GFP).

### Preparation of alginate hydrogels

All alginate gels were initially prepared at a 3% (w/v) concentration, for ultimate dilution to 2% after loading them with therapeutic. A 3% alginate mixture (w/v) was made with alginic acid powder (Sigma-Aldrich) and alpha Minimum Essential Medium (αMEM, Gibco) then left overnight at room temperature. This solution was sterile filtered (0.2 μm, Pall) and transferred into 1mL syringes. Therapeutics were then added directly to the alginate. The crosslinker, calcium sulfate (0.21g CaSO_4_ / mL distilled H_2_O) was loaded into a separate 1mL syringe and was mixed vigorously with the alginate solution for approximately one minute. Hydrogels were kept at 4°C or on ice until use.

### Synthesis of CRISPR-Cas9 modified bacteriophages

*S. aureus* strain RF122 harboring CRISPR-Cas9 modified bacteriophage was cultured in BHI broth to the mid-exponential phase (OD600 at 0.3).[21] To induce CRISPR-Cas9 modified bacteriophage, mitomycin C (1 μg/mL, Sigma-Aldrich) was added to the culture and further incubated at 30°C with shaking at 80 RPM. A complete lysis of culture typically occurred within 2-3 hours. The clear lysate was sterilized with syringe filers (0.22 μm, Nalgene). The concentration of phage was calculated by determining the plaque-forming units using a soft agar (0.5%, w/v) overlaying method.[21]

### Kirby-Bauer analyses

To analyze the bactericidal activity of therapeutics, a Kirby-Bauer assay was performed as previously described, with slight modifications.[23] Stock solutions of fosfomycin (50 mg/mL) and phage (~10MOI/mL) were prepared in phosphate buffered saline (PBS). Using these stock solutions, a total of 10μL of: (i) fosfomycin, (ii) phage, (iii) dual: fosfomycin (5μL) and phage (5 μL), or (iv) PBS alone were directly applied to bacterial lawns, without the use of disks as traditionally described. The applied solutions were allowed to set undisturbed for approximately 5-10 minutes at room temperature, and were then incubated at 37°C for 24h. The zones of inhibition were then measured and recorded.

### Qualitative and quantitative bactericidal activity on biofilms

For qualitative *in vitro* evaluation of antibiofilm efficacy, a 6-well tissue culture plate was pre-coated with 2% human serum for 24 hours, after which *Staphylococcus aureus* ATCC 6538-GFP was cultured in tryptic soy broth (TSB) supplemented with 2% glucose for 72 hours. After gentle washing with PBS, TSB supplemented with vancomycin (256, 512, or 1024 μg/mL), fosfomycin (16, 64, 128 μg/mL) or bacteriophage (5, 10, or 25 multiplicity of infection (MOI)) was added to the biofilm and incubated for 24 hours. After gentle washing with PBS three times, remaining biofilm indicated by GFP signal was measured using Cytation 5 plate reader (BioTek).

To quantify the antibiofilm activity of selected therapeutics delivered by alginate hydrogels, fosfomycin, phage, or dual therapeutic was loaded in 2% alginate hydrogel and overlaid on top of the biofilms. As a control, empty 2% alginate hydrogel was used. BHI broth was added and cultures were incubated at 37°C for 24 h. After removing BHI broth, the entire 2% alginate hydrogel and biofilm were harvested and vigorously washed with PBS by centrifugation to remove residual therapeutic. A serial dilution in PBS was plated onto BHI plates to determine viable bacterial counts.

### Rat osteomyelitis model

All procedures (Fig. 1) were performed in accordance with the Institutional Animal Care and Use Committee (IACUC) of Mississippi State University. Charles River Sprague Dawley female rats, 13 weeks old, were housed with 12h light/dark cycles and were provided food and water *ad libitum*. Rats were administered slow release buprenorphine (1.0-1.2 mg/kg BW, ZooPharm) pre-operatively for pain relief. Rats were anesthetized with isoflurane at an initial concentration of 2-3%, and maintained at 1-2%. After sterile preparation of the left hindlimb by fur removal and alcohol and chlorhexidine scrubs, the skin was incised with an anterior approach, from the level of mid-diaphysis to the patella, along the lower half of the femur. The muscle tissue was separated using blunt dissection along the muscle bundle divisions on the anterolateral side of the femur. In the mid-diaphysis, a 1.2 mm (diameter) bicortical defect was created with a pneumatic drill (Conmed Hall), and a #65 drill bit (McMaster-Carr). To mimic contamination of orthopedic screws with *S. aureus* occurring in development of osteomyelitis *in vivo*, sterile orthopedic screws (Antrin Miniature Specialties, #00-90) were placed into 200 μL of a bacterial suspension (~1×10^8^ CFU) of ATCC 6538-GFP for approximately 5-10 minutes (average 6.5 min). The screw was then placed into a 96-well plate to dry for up to 6 minutes (average 4 min). The bacterial load of these contaminated screws was approximately 5×10^4^ CFU, determined by placing screws into 1mL of PBS, vigorously vortexing to elute bacteria from the screw, then serially diluting the eluents for bacterial counting on BHI agar plates. *In vitro* characterization of bacterial load based on (i) the time screws remained in culture (soak time) and (ii) dry time was performed using the same procedure as for bacterial counts from *ex vivo* screws. To assess the effect of soak time, dry time was kept at a constant 5 minutes, and similarly for the effect of dry time, soak time was kept constant at 5 minutes. To complete the *in vivo* procedure, the superficial fascia lata and skin were closed with sutures. Longitudinal monitoring of infection at days 1, 3, and 6 post-infection was performed via radiographs with fluorescent overlays using the IVIS Lumina XRMS II system (PerkinElmer).

**Fig 1.**
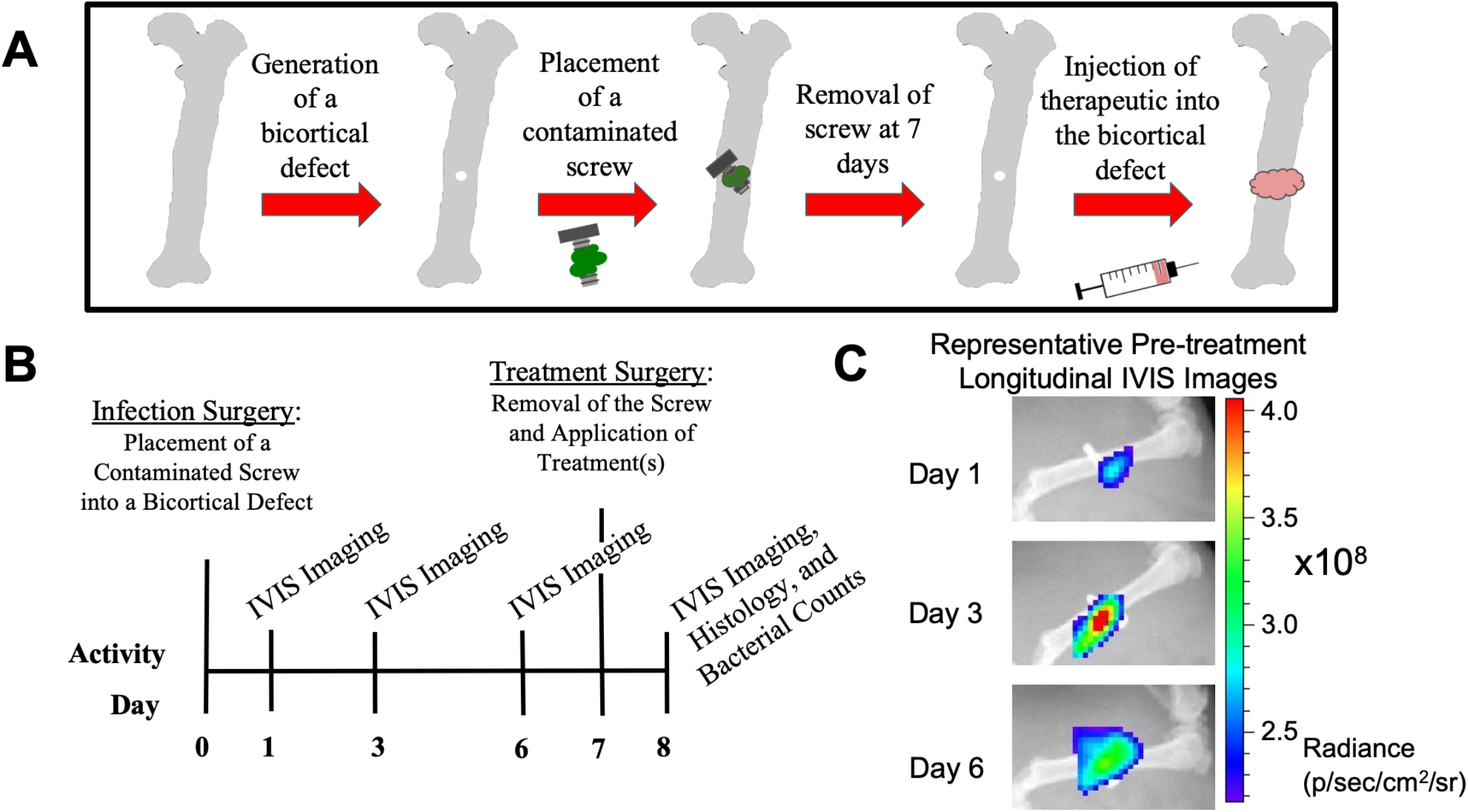
Overview of *in vivo* experimental procedure. (A-B) On day 0 (infection surgery day), a bicortical defect (drill hole) was generated in the mid-diaphysis of the left femur. A contaminated orthopedic screw was then fastened into this space, and left for 7 days to generate robust osteomyelitis and soft tissue infection. At day 7, the orthopedic screw was removed and 100μL of therapeutic(s) were injected into the defect space. At day 8, 24h after treatment, soft tissues and bone samples were collected for histology, scanning electron microscopy, and bacterial counts. (C) At days 1, 3, and 6, IVIS imaging was performed to track infection progression.

After a 7 day infection period, the area was accessed along the original incision line. The infected screw was removed and placed into 1mL PBS or fixative for bacterial counting or SEM, respectively. Then, 100 μL of fosfomycin (3 mg), phage (MOI 3), dual (3 mg fosfomycin and MOI 3 phage), or PBS loaded into 2% alginate hydrogel was injected into the lateral end of the bicortical defect, with excess hydrogel pooling in the medial, underlying soft tissue.

On day 8, approximately 24 hours post-treatment, animals were sacrificed via CO_2_ inhalation. The hindlimb was initially cleaned with chlorhexidine, and sterile instruments were used to disarticulate the femur and adjacent soft tissues for further evaluation. For bacterial counting, bone samples were initially minced using sterile bone rongeurs and further processed using a homogenizer (Cole-Parmer, LabGEN7, 30s at setting 2-3, 30s at setting 9-10). Soft tissue samples were minced using sterile surgical scissors, then homogenized (30s at setting 2-3, 30s at setting 7-8). Following initial processing, homogenates were vortexed (2000 RPM, 1 minute), diluted as necessary, spread onto BHI agar plates, and incubated for 24 hours at 37°C for enumeration, with a detection limit set at 25-250 colonies.

### Electron microscopy and histological analyses

Screw samples for electron microscopy analysis were collected during revision surgeries on day 7 immediately prior to application of treatment, and placed directly into a fixative consisting of 5% glutaraldehyde, 2% paraformaldehyde (w/w) in a sodium cacodylate buffer (a modified “Karnovsky’s” solution).[24] For electron microscopy analysis of infected bone, a representative control femur (empty alginate group) was collected at day 8, broken along the screw line with sterile bone rongeurs, and immediately placed in fixative for 24h. For both the screw and bone samples, no dehydration series was performed in order to preserve the biofilm. Prior to imaging, both samples were placed onto stubs with carbon tape and sputter coated (Quorom Tech Model # SC7640) with platinum at 30 mA and 3.5 kV for 3-5 minutes. All samples were then imaged using FESEM (Carl Zeiss AG-SUPRA 40).

For histological analyses, the infected femur and adjacent soft tissues were placed into 10% formalin for 48h, at ~20°C. Bone samples were decalcified for 5 days in Kristensen’s solution, then rinsed and placed into 10% formalin.[25] Tissues were routinely processed, embedded in paraffin, sectioned at 5μm, and Gram or hematoxylin and eosin (H&E) stained.

### Statistical analyses

All statistical analyses were performed using either GraphPad Prism 8 or SAS software systems. For the Kirby-Bauer assay, a one-way analysis of variance (ANOVA) with Tukey’s multiple comparisons test was performed. For the *in vitro* antibiofilm assay, a two-way ANOVA with Tukey’s multiple comparisons were performed. For bacterial counts from directly prepared and *ex vivo* orthopedic screws, a one-way ANOVA was performed, with Sidak’s multiple comparisons. All aforementioned statistical analyses were performed using GraphPad Prism 8 (GraphPad Software, Inc.). For bone and soft tissue *ex vivo* bacterial counts, general linear models using PROC MIXED in SAS (SAS Institute, Inc.) were performed, with pairwise comparisons using Tukey’s (comparing treatment groups to one another) or Dunnett’s (comparing each treatment to control) tests. An alpha level of 0.05 was used to determine statistical significance for all methods. Data are presented as mean ± standard deviation (SD).

## Results

### Integration of GFP into *S. aureus*

Longitudinal analyses of infection progression and regression is an ideal tool for new model generation. Plasmids harboring reporter genes such as luminescence and fluorescence have been most commonly used; however, it is necessary to constantly provide antibiotic selective pressure to prevent a loss of plasmid, which is not achievable with *in vivo* models. In this study, we integrated the GFP reporter gene into the genome of *S. aureus* ATCC 6538 strain for the stable and accurate assessment of bacterial growth. All *S. aureus* ATCC 6538-GFP colonies grown in BHI plates without antibiotic selective pressure were highly fluorogenic over a span of 7 days (Fig. 2A). Furthermore, ATCC 6538-GFP recovered from an *ex vivo* orthopedic screw at day 7 post-infection was still highly fluorogenic (Fig. 2B). These results demonstrated that *S. aureus* ATCC 6538 chromosomally integrated with GFP can be used for real-time monitoring of bacterial proliferation *in vitro* and *in vivo*.

**Fig 2.**
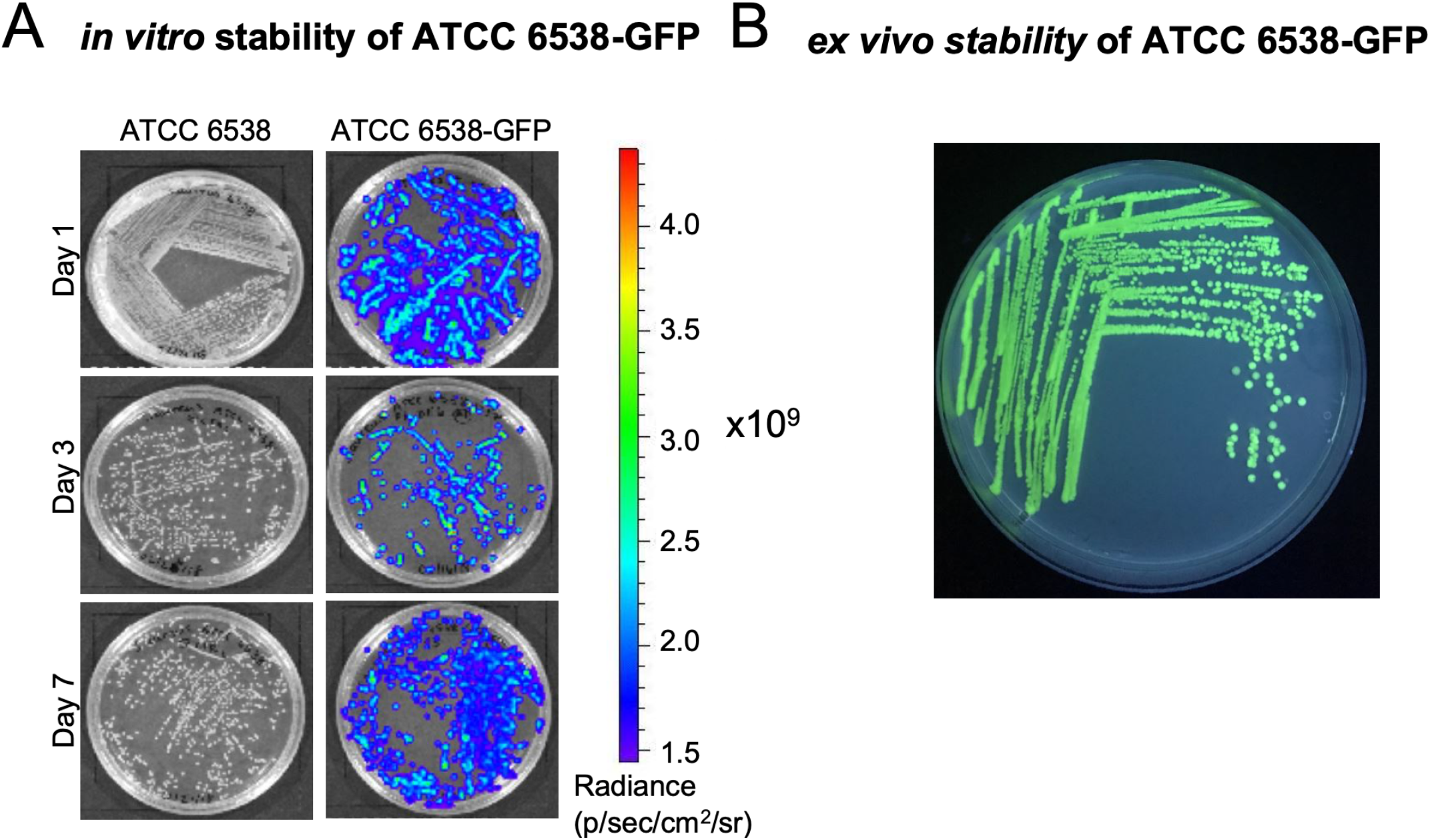
Integration of GFP into ATCC 6538. (A) Phenotypic expression confirming integration of GFP into ATCC 6538 was stable over a 7 day period. (B) A *Staphylococcus aureus* CFU isolated from a contaminated screw *ex vivo* at day 7 revealed GFP expression continued after growth over a week *in vivo*.

### Kirby-Bauer analyses

For initial investigation of selected therapeutics, a Kirby-Bauer assay was performed (Fig. 3A). All therapeutics—fosfomycin, phage, and dual—had a larger zone of inhibition than the PBS control (“a”, p<0.0001), which generated no zone of inhibition (“N.D.”) Dual and fosfomycin also resulted in a zone of inhibition greater than phage treatment (“b,” p<0.0001).

**Fig 3.**
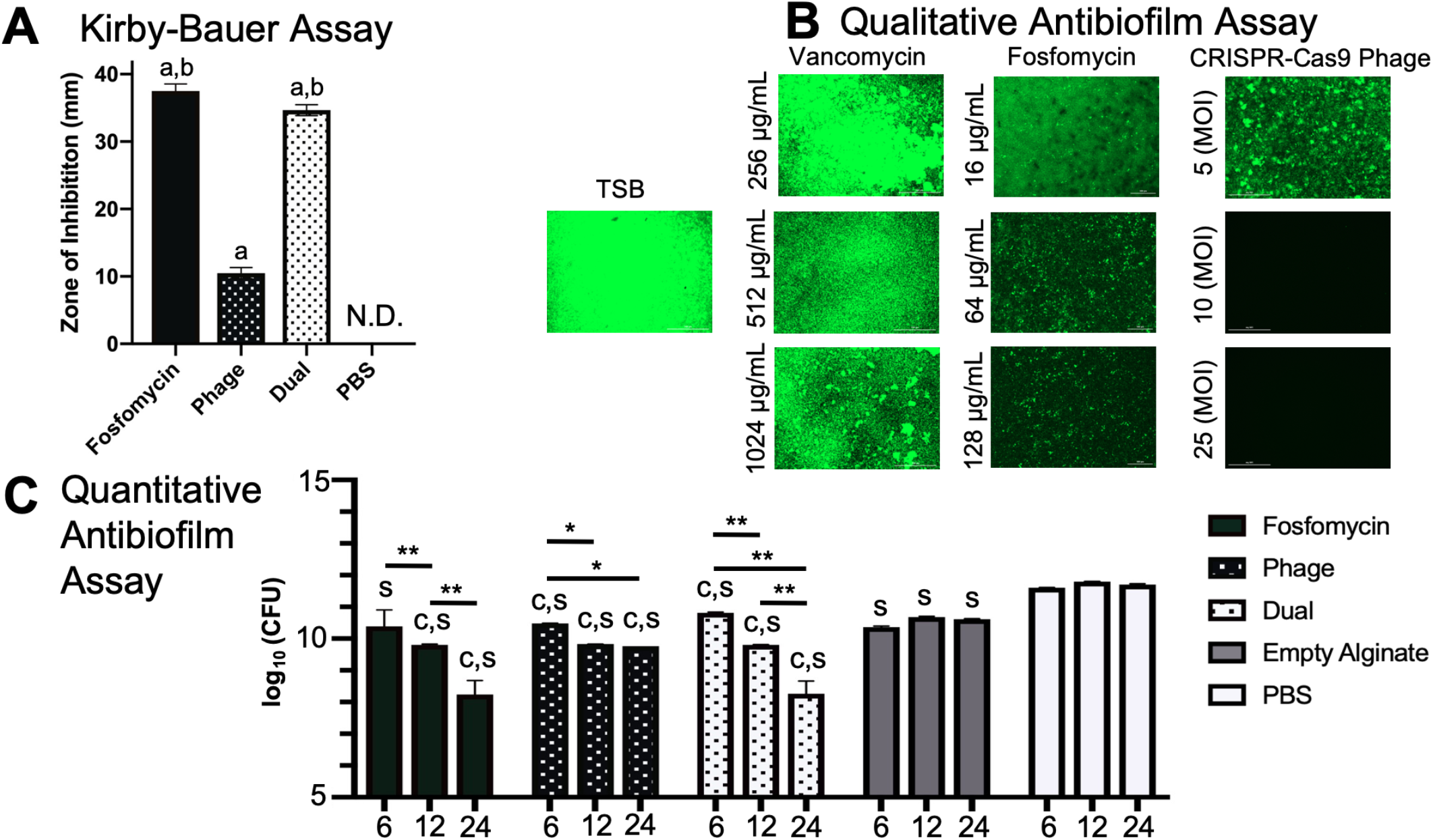
*in vitro* Analyses of Therapeutic Bactericidal Activity. (A) From the Kirby-Bauer assay, all therapeutics delivered via PBS had a greater antibacterial effect than the PBS control (“a”, p<0.0001), which generated no zone of inhibition (“N.D.”). Dual and fosfomycin therapeutics also generated a zone of inhibition greater than phage treatment (“b,” p<0.0001). (B) Vancomycin (256-1024 μg/mL), fosfomycin, (16-128 μg/mL) and bacteriophage (MOI 5-25) delivered via PBS revealed varied bactericidal activity, where green indicated bacterial vitality and black indicated a lack of bacterial presence. Interestingly, vancomycin appeared to have little to no efficacy on biofilms at all concentrations. Fosfomycin, in contrast, showed efficacy at 64 and 128 μg/mL, a dose range approximately one-tenth the vancomycin doses utilized. Bacteriophage was effective at an MOI of 10, indicated by the black panel revealing no viable *S. aureus*. (C) Compared to the empty alginate group, alginate-loaded fosfomycin, phage, and dual therapeutic-treated biofilms had lower bacterial loads at 6, 12, and 24 hours, except the fosfomycin group at 6 hours (“c,” p<0.05). Interestingly, all groups (fosfomycin, phage, dual, and empty alginate gel) had lower growth at all time points compared to the PBS control, i.e. the empty alginate gel exerted a killing effect (“s,” p<0.05). Within individual treatment groups over time, increased antimicrobial effects were observed. The fosfomycin treated biofilms were different at 6 and 12 hours, and at 12 and 24 hours (**p<0.0001). Phage treated groups were different at 6 and 12 hours, and at 6 and 24 hours (*p<0.05). Dual treated biofilms were different at 6 and 12 hours, at 6 and 24 hours, and at 12 and 24h (**p<0.0001). The empty alginate and PBS controls resulted in no changes over time.

### Qualitative and quantitative bactericidal activity on biofilms

Biofilms are generally considered the greatest agent of osteomyelitis treatment failure, and thus are important to consider when developing therapeutics. For this reason, it is important to evaluate the efficacy of novel therapeutics on robust biofilms, for translation into relevant clinical scenarios. Antibiofilm efficacy of therapeutics *in vitro* was characterized utilizing two different analyses: (i) qualitative fluorescent (phenotypic) assessment and (ii) quantitative bacterial counting. Surprisingly, vancomycin, one of the most commonly utilized antibiotics for difficult osteomyelitis cases, appeared to have little or no effect on biofilms, as indicated by the presence of green fluorescing *S. aureus* (Fig. 3B). Fosfomycin is a small molecular weight (138 g/mol) broad-spectrum antibiotic and promising therapeutic option against biofilm.[26] Here, fosfomycin appeared to remove biofilm at 64 and 128 μg/mL, doses much lower than vancomycin. The CRISPR-Cas9 modified bacteriophage has dual killing mechanisms: (i) a direct lysis of target bacteria by holin or murein hydrolase, and (ii) CRISPR-Cas9 nuclease activity. From qualitative fluorescent analyses, it was determined than a phage MOI of ~10 was effective in clearing biofilm infection (Fig. 3B).

Alginate is a versatile biopolymer used for prolonged, localized availability of therapeutic.[27] Antibiofilm assays with bacterial counts were utilized to qualitatively assess the effects over time of selected therapeutics delivered via alginate (Fig. 3C). Compared to the empty alginate group, fosfomycin, phage, and dual therapeutic-treated biofilms had significantly lower bacterial loads at 6, 12, and 24 hours, except the fosfomycin group at 6 hours (“c,” p<0.05). Interestingly, all groups (fosfomycin, phage, dual, and empty alginate gel) were significantly lower at all time points compared to the PBS treatment, i.e. the empty alginate gel exerted a killing effect (“s”, p<0.05). In all treatment groups where alginate was loaded with therapeutic(s), antibiofilm effects increased over time. The fosfomycin treated biofilms were different at 6 and 12 hours, and at 12 and 24 hours (**p<0.0001). Phage treated groups were different at 6 and 12 hours, and at 6 and 24 hours (*p<0.05). Dual treated biofilms were different at 6 and 12 hours, at 6 and 24 hours, and at 12 and 24h (**p<0.0001). As expected, the empty alginate and PBS controls did not change over time.

### Bacterial load on orthopedic screws

To generate consistent infection in the osteomyelitis model, contaminated orthopedic screw preparation had to first be characterized. Two parameters of screw preparation were evaluated: soak time and dry time. Soak time (5-20 min) of screws appeared to increase somewhat proportionally with respect to bacterial load (Fig. 4A). Dry times from 0-10 min appeared to have little effect on bacterial load; at 20 min, decreased viability of *S. aureus* was observed (Fig. 4B).

**Fig 4.**
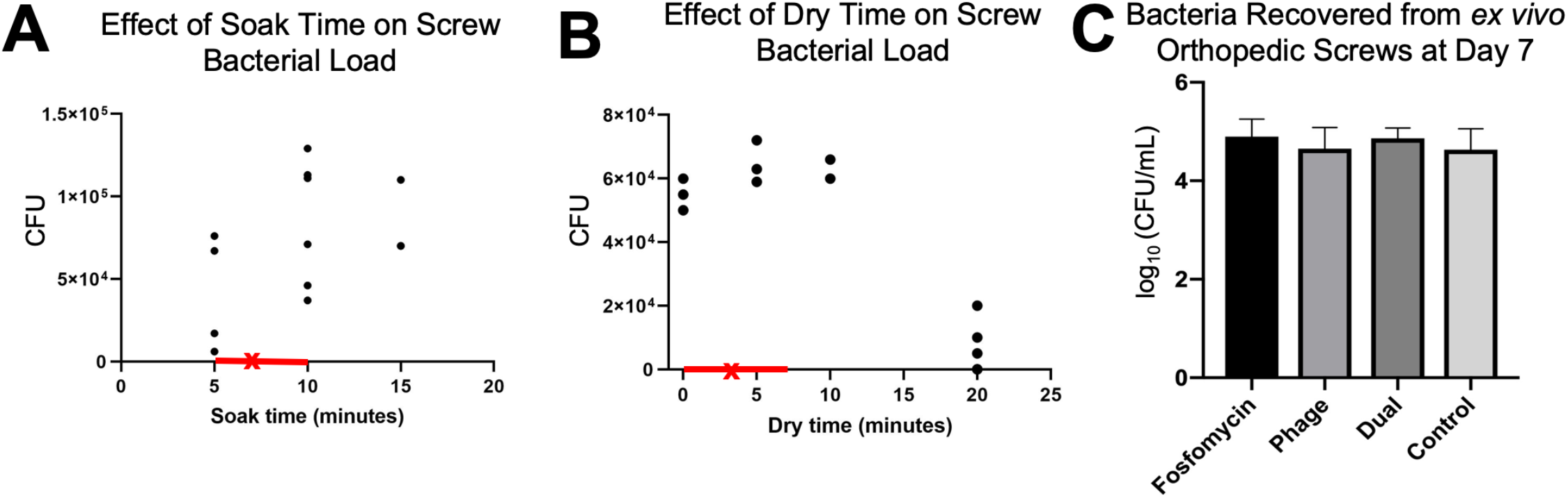
Effect of soak time and dry time on bacterial load of orthopedic screws. (A) Soak time of sterile orthopedic screws in *Staphylococcus aureus* appeared to linearly relate to ultimate bacterial load. (B) Dry time did not appear to decrease ultimate bacterial load, until 20 minutes of dry time. (A-B) The ranges of soak time and dry time used for preparation of screws for the *in vivo* study are indicated by the red portions of the x-axes. The average time for soak and dry times for orthopedic screws used *in vivo* is marked by an “x”. (C) Bacterial counts collected from orthopedic screws at day 7 *ex vivo* indicated a similar ultimate bacterial load among what would become different treatment groups.

In the *in vivo* model, orthopedic screws removed at day 7 to allow injection of therapeutic into the infected defect space were analyzed for bacterial load to confirm all treatment groups began with a similar extent of infection. Bacterial counts from *ex vivo* screws indicated similarly severe infection among all samples (treated immediately following), with an average 8.19×10^4^ CFU/mL bacterial load (Fig. 4C). Per what would become individual treatment groups, calculated averages were: 1.05×10^5^, 7.50×10^4^, 8.20×10^4^, and 6.57×10^4^ for fosfomycin, phage, dual, and control groups, respectively. No significant differences between any groups were observed.

### Scanning Electron Microscopy

Representative SEM images of screws collected at day 7 post-infection revealed an abundance of gram positive cocci (*S. aureus*), as expected, along the distal portion of the screw and within the ridges of the screw throughout its length (Fig. 5A-B). In bone fragments collected along the screw line (defect site) of an untreated (empty alginate) control sample at day 8 (24h post-treatment), gram positive cocci, presumably *S. aureus*, were visible. (Fig. 5C-D).

**Fig 5.**
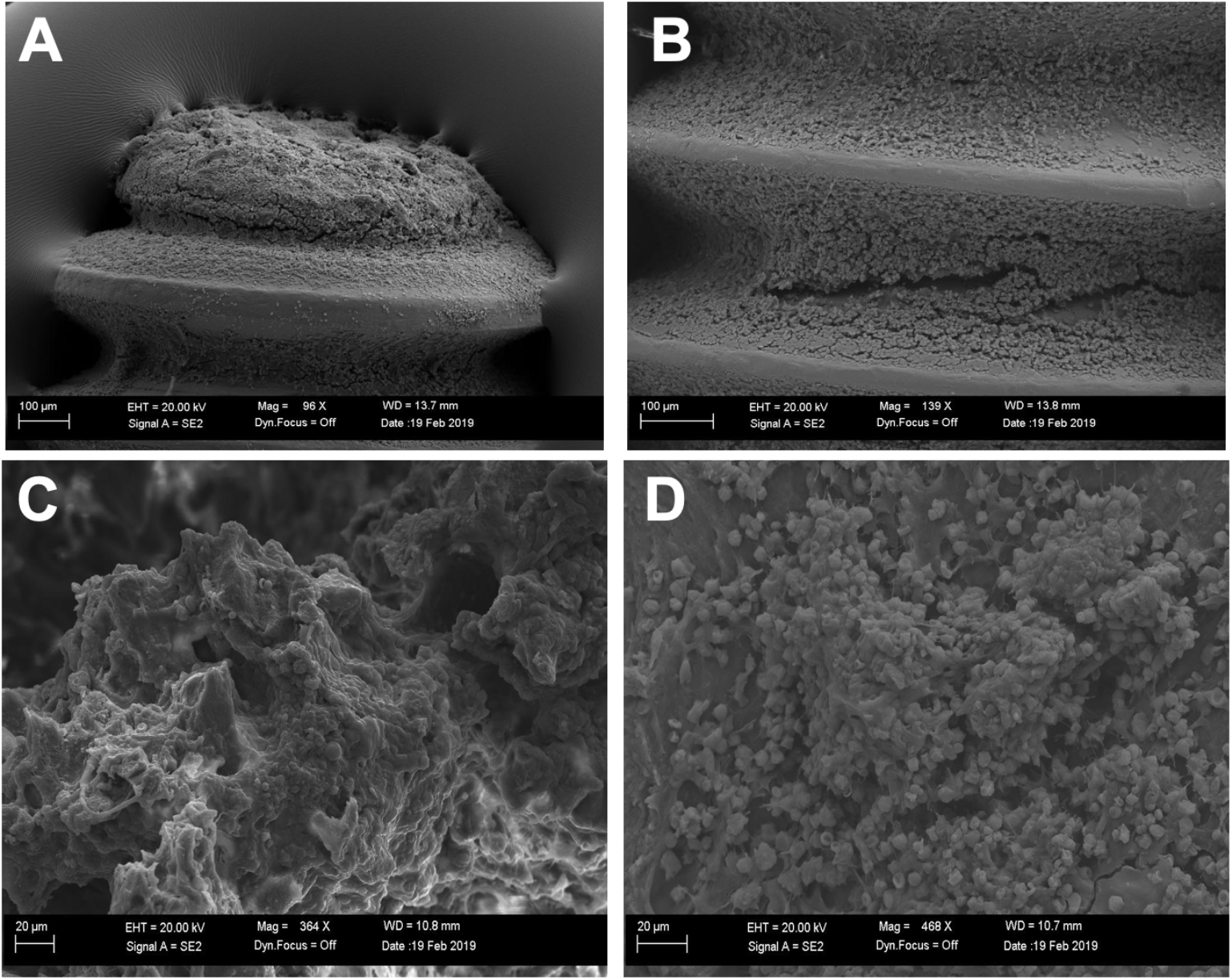
Bacteria on *ex vivo* orthopedic screws and bone. (A-B) Gram positive cocci, and what appears to be biofilm, was evident on the distal portion (A) and between threads (B) of an *ex vivo* orthopedic screw excised at day 7. (C-D). Femur sample adjacent to the defect/screw site collected at day 8 from untreated (empty alginate) control revealed dispersed gram positive cocci.

### Bacterial load within bone and soft tissue

The average bone bacterial counts per treatment group were as follows: (i) control: 4.197 ± 0.289, (ii) fosfomycin: 3.401 ± 0.924, (iii) phage: 4.076 ± 0.268, and (iv) dual: 3.607 ± 0.316 (Log_10_(CFU)), Fig. 6A). Fosfomycin bacterial counts were lower than empty alginate control (p=0.0083) and phage (p=0.0486).

**Fig 6.**
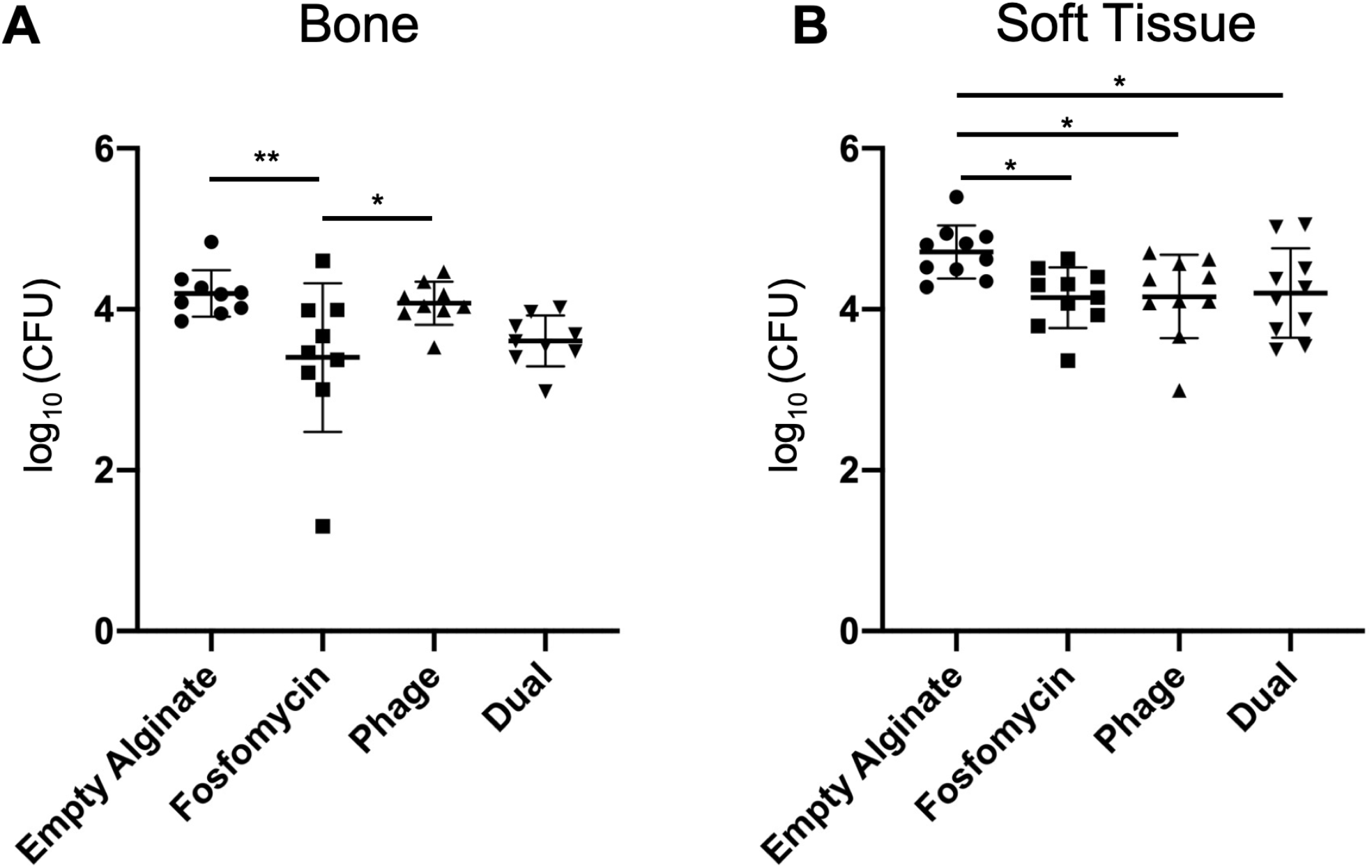
Bacterial counts from bone and soft tissue. (A) Bacterial counts from bone tissue harvested from fosfomycin treated animals were lower than those for empty alginate (**p=0.0083) and phage (*p=0.0486) groups. (B) In soft tissue, bacterial loads were reduced in all three treatment groups—fosfomycin (*p=0.0225), phage (*p=0.0265), and dual (*p=0.0430)—compared to empty alginate.

The average soft tissue bacterial counts per treatment group were as follows: (i) control: 4.713 ± 0.331, (ii) fosfomycin: 4.146 ± 0.377, (iii) phage: 4.160 ± 0.516, and (iv) dual: 4.201 ± 0.556 (Log_10_(CFU)), Fig. 6B). Bacterial counts were lower in fosfomycin (p=0.0225), phage (0.0265), and dual (p=0.0430) treated groups compared to empty alginate control.

### Histology

Bone samples stained with H&E or Gram revealed strong evidence for development of a severe osteomyelitis infection. The areas at the site of the screw were characterized by extensive remodeling within the medullary cavity, with replacement of marrow cells by a central area of neutrophils surrounded by fibrovascular proliferation and reactive bone. Within the cortex, at the site of screw placement, was mild bone necrosis characterized by empty lacunae and bone loss. Along the periosteal surface there was locally extensive proliferation of woven bone (periosteal proliferation). Additionally, abundant gram-positive cocci were localized within the bone. No differences in the extent of infection among any groups were apparent. The outcome of the fosfomycin-treated bone histology varied from the other samples, due to sample damage during processing. Ultimately, this prevented the collection of images along the screw line (where the majority of bacteria and inflammation were localized), as seen with the other groups. However, the proliferation of woven bone, as a reactive process on the cortical surface, is visible (Fig. 7B). Within the phage-treated bone sample, a linear track of gram-positive bacteria in the bone at the original site of the screw line was visible (Fig. 7C). In the dual-treated sample, neutrophilic inflammation surrounded by reactive bone and fibrosis was observed (Fig.7D left, *).

**Fig 7.**
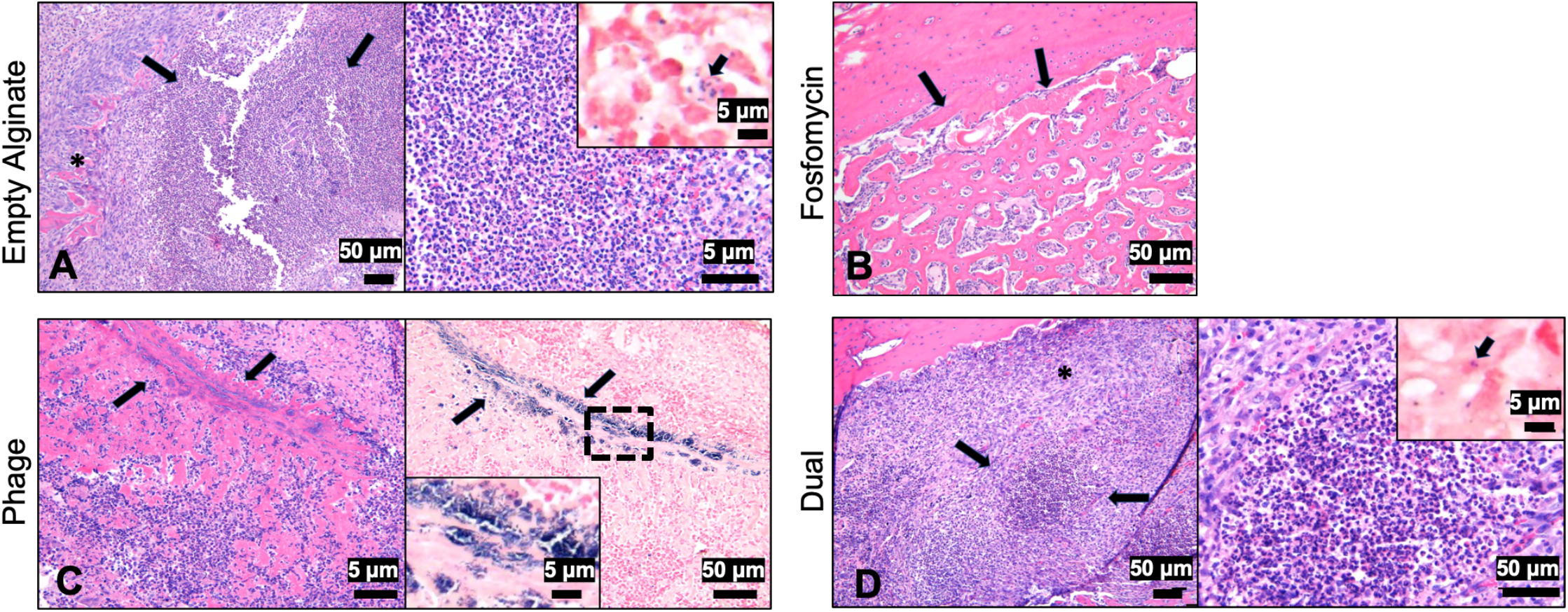
Histology of infected bones one day post-treatment. (A, left) At the site of the screw of the empty alginate control was marked neutrophilic inflammation (arrows) with bone loss, surrounded by reactive bone and fibrosis (*, bar=50μm). (A, right) A higher magnification of the area of bone loss with large numbers of neutrophils (bar=5μm). Inset demonstrating gram positive bacteria within and outside macrophages (arrow, bar=5μm). (B) Within the fosfomycin treated bone, periosteal proliferation of woven bone was noted (arrows, bar=50μm). (C, left) A linear track in the bone at the site of the screw (arrows) with abundant neutrophils and reactive bone and fibrosis was observed (arrows, H&E, bar=5μm). (C, right) Gram staining demonstrating aggregates of basophilic bacteria along the screw site (bar=50μm). Inset is a higher magnification of the bacteria (bar=5μm). (D, left) At the site of the screw was marked neutrophilic inflammation with bone loss (arrows), surrounded by reactive bone and fibrosis (*, bar=50μm). (D, right) A higher magnification of the area of bone loss with large numbers of neutrophils (bar=50μm). Inset demonstrating gram positive bacteria within and outside macrophages (arrow, bar=5μm).

## Discussion

In this manuscript, a previously developed CRISPR-Cas9 modified bacteriophage therapeutic, which was successful in treating external dermal infection[21], was evaluated as a therapeutic for internal osteomyelitis and contiguous soft tissue infection in a rat model using a biofilm forming strain of *S. aureus*. For real time monitoring of *S. aureus*, GFP was chromosomally integrated into *S. aureus* ATCC 6538 strain by homologous recombination. We demonstrated that *S. aureus* ATCC 6538 strain carrying the GFP gene stably maintained the GFP phenotype without antibiotic selective pressure *in vitro* and *in vivo*.

The therapeutic effects of vancomycin, fosfomycin, CRISPR-Cas9 modified bacteriophage, and fosfomycin-bacteriophage dual treatments were evaluated against biofilm *in vitro*. Vancomycin is considered a last resort treatment for *S. aureus* infection. Recent guidelines recommend vancomycin trough concentrations between 15 and 20 μg/mL for effectively controlling *S. aureus* infection.[28] As biofilms are typically more difficult to treat than planktonic bacteria, a much higher concentration of vancomycin (256-1024 μg/mL) was used here. Although the planktonic *S. aureus* ATCC 6538 strain was sensitive to vancomycin at 2 μg/mL within BHI broth (data not shown), in biofilm form, it was highly resistant to vancomycin even at 1024 μg/mL. Biofilms consist of a group of bacteria and their byproducts such as extracellular polymeric substances (EPS) including proteins, DNA, RNA, polysaccharides, and peptidoglycans. These EPS materials provide physical barriers to penetration of antibiotics to the inner viable population of bacteria in the biofilm. Vancomycin, a large glycopeptide antibiotic with a molecular weight of 1,449 g/mol, binds to the D-ala-D-ala terminal amino acid at the stem of pentapeptide crosslinking peptidoglycan for efficacy[29]. Thus, resistance to vancomycin by biofilm may be explained by poor penetration of vancomycin due to its bulky size, which could have led to entrapment at the peptidoglycan layer of biofilm. In contrast, fosfomycin showed better efficacy against biofilm at much lower (64 and 128 μg/mL) doses. Fosfomycin is a small bactericidal antibiotic with a molecular weight of 138 g/mol. It interferes with the first step of peptidoglycan synthesis by inhibiting the phosphoenolpyruvate synthetase[30]. Thus, the enhanced fosfomycin efficacy could be explained by better penetration of fosfomycin due to its small size and its inhibition of the first step of peptidoglycan synthesis. From qualitative fluorescent analyses, it was determined than a phage MOI of ~10 was effective in clearing biofilm infection. This is similar and in some cases an improvement upon *in vitro* evaluation of phage treatments discussed in literature, with biofilm eradication reported with MOI 10-100.[21,31,32] Alginate hydrogel served as an effective delivery vehicle, enabling increasing effects against biofilm over a 24h period, for fosfomycin, phage, and dual treatments *in vitro*. Previously, we have observed similar sustained effects of bone morphogenetic protein-2 released from and retained within alginate hydrogels *in vitro* and *in vivo*.[18,27]

For further evaluation of the CRISPR-Cas9 phage, motivated by augmented *in vitro* efficacy relative to antibiotic controls, we developed a clinically relevant model of implant-related osteomyelitis. In human cases of osteomyelitis, chronic infection is diagnosed after a 6-week period of infection, while our model had only a 1-week infection period. Based on SEM images, 1 week appeared to be sufficient to induce severe infection, including biofilm, in this rat model. By culturing *S. aureus* on orthopedic screws, infection was localized to the femur and surrounding soft tissue, as indicated by fluorescent imaging and histology. Fluorescent imaging served as a qualitative tool for longitudinal infection progression/regression, although no direct correlation between radiance output and bacterial load were observed (data not shown). This could be attributed at least in part to a residual GFP signal that likely exists after bacterial cell death, due to the persistence of the GFP. Clinically, debridement accompanied by long-term antibiotic administration is the gold standard for osteomyelitis treatment.[33] In this study, we have avoided debridement altogether so as to limit potential clearing of infection from any source other than the therapeutics delivered. For future studies, debridement may be included to more readily mimic the clinical scenario and enable evaluation of larger antibacterial materials such as scaffolds or putties.

From bacterial counts performed on excised soft tissues, it was determined that severe soft tissue infection accompanied the expected high bacterial load in the bone samples. In clinical cases of osteomyelitis, soft tissue infection is a common pathological finding of osteomyelitis infection progression[34–36]. In this model, soft tissue infection likely developed due to the distal end of the orthopedic screw resting freely within the soft tissue medial to the defect site. On excised orthopedic screws collected on day 7, scanning electron microscopy indicated a purulent, thick biofilm layer of *Staphylococci* on the end of the orthopedic screw. Based on *in vitro* results, the process of orthopedic screw preparation can be used to tailor the extent of infection, as soaking the screws for a shorter period of time would be expected to introduce less *S. aureus* into the bone and as a result induce a less severe infection. Histology results support the development of severe osteomyelitis infection, with disease hallmarks such as neutrophilic inflammation, reactive bone, fibrosis, and gram-positive bacteria. Within the 24-hour time frame of this study, no differences among treatment groups were apparent. If later time points were evaluated, the differences noted in bacterial counting would likely be more readily reflected histologically.

Although only the fosfomycin group resulted in reduced bacterial load in the femur, in soft tissue, all three treatments, including phage alone and phage with fosfomycin (dual) led to lower bacterial counts compared to empty alginate. It should be noted than an extremely high dose of fosfomycin (3g) was administered to the rat femur in this study. In humans, a 3g oral dose is recommended for treatment of urinary tract infections.[26] Conversely for bacteriophage dose, although a minimum effective MOI of ~10 was observed *in vitro*, *in vivo* only MOI of ~3 was able to be delivered due to: (i) the volume of alginate hydrogel delivered to the small defect site (100 μL total, but only ~10 μL fit into the defect itself), and (ii) the thicker consistency of phage solution, limiting the concentration that could be prepared in alginate hydrogel. Collectively, these discrepancies in dosing likely limited the efficacy of phage treatment alone in osteomyelitis mitigation relative to antibiotic with and without phage. Furthermore, the canaliculi of cortical bone may provide an ideal environment for bacteria to evade treatment.[37] It should also be noted that in this study, a one-time 100 μL treatment was applied locally. Clinically, osteomyelitis is treated via debridement and systemic antibiotics for several weeks.[33] Similarly, success with bacteriophage therapy has been associated with continuous, prolonged delivery of the virus. A bacteriophage cocktail was used to successfully clear femoral infection with four intraperitoneal doses of phage (100 μL of ~2×10^12^ plaque forming units (PFU)/mL) over the span of 48 hours.[38] Another study adopted a treatment regimen for tibial osteomyelitis consisting of a once daily 3×10^8^ PFU/mL intramuscular bacteriophage injection for 14 days, which resolved the infection.[39] Recently, a case report was published describing the success of a weekly injection of bacteriophage over a seven-week period for human digital osteomyelitis.[36] Collectively, these data suggest that sustained, localized concentrations of phage may be necessary for efficacy in treatment of bone infection. In future studies, a greater initial dose of phage therapeutic should be considered, or a longer duration of treatment achieved with a delivery vehicle capable of tailored release of therapeutic. Given a higher phage dose and/or prolonged availability, it is possible that the efficacy of phage observed here *in vitro* could be matched *in vivo*. Furthermore, it may be more advantageous to use alternating doses of the antibiotic and phage therapeutic over time, rather than a combined simultaneous application.[40,41] In this study, no additive effect of fosfomycin or phage was observed in the dual treatment group. Only a 24h treatment period was evaluated, which may have limited the effect of our selected therapeutics, as later time points may have allowed therapeutics, especially those containing phages (which must replicate for optimal bactericidal activity), to have a cumulative effect. Nonetheless, our challenging composite tissue infection model enabled efficient, rapid testing of antimicrobial therapeutics using a biofilm forming strain of *S. aureus*.

Despite the prevalence and severity of osteomyelitis, no bacteriophage-based treatment for the disease has reached clinical trials in the United States. As populations of MDR-bacteria continue to spread and new strains are isolated, engineering novel therapeutics will be essential to augment the dwindling list of effective, available antibiotics. Phage therapy has great potential to fill this niche, as phages can be made readily and at a low cost. Using CRISPR-Cas9 technology as in this study, phages can be modified to have a wide host range.[21] By contributing to the pipeline of bacteriophage therapeutic evaluation compared to traditional antibiotics, the goal of this work was to demonstrate efficacy of phage against bone and soft tissue infection. Enhanced bactericidal activity of CRISPR-Cas9 phage was demonstrated *in vitro* against biofilm, when compared to conventionally used vancomycin and fosfomycin antibiotics. Subsequently, an implant-related model of osteomyelitis and soft tissue infection was established, confirmed with histological, radiographic, and scanning electron microscopy analyses. Although phage did not mitigate bone infection 24h post-treatment, soft tissue infection was reduced 24h following treatment with bacteriophage, and notably to the same extent as treatment with high dose antibiotic. To improve bacteriological outcomes in the future, further investigations of delivery vehicles and optimal dosing are warranted.

## Acknowledgements

This research was supported by the NIH Center of Biomedical Research Excellence in Pathogen-Host Interactions (2P20GM103646) and the Office of Research and Economic Development at Mississippi State University. The authors would like to thank Dr. Jean Feugang, Dr. Seongbin Park, and the USDA-ARS Biophotonics Initiative (58-6402-3-018) for supporting our use of the IVIS Lumina XR system, Dr. Robert Wills for statistical analyses, and Dr. Lucy Senter and Dr. Bridget Willeford for veterinary care. Thanks also to Jamie Walker, Delisa Pennell, Dr. Hayley Gallaher, Weitong Chen, Kali Sebastian, Kristen Lacy, Christine Grant, Drew Moran, Ryan Yingling, Alex Feaster, Hannah Bostick, Christina Moffett, Sonja Jensen, Anna Marie Dulaney, and Luke Tucker for assistance with surgeries.

